# Brain–AI Alignment in Naturalistic Movies

**DOI:** 10.64898/2025.12.03.692164

**Authors:** Muwei Li

## Abstract

Naturalistic paradigms offer a powerful window into human cognition, but it remains difficult to link rich, continuous movie content to distributed brain activity in an interpretable way. In this study, I use a multimodal large language model (Gemini) as an automated “semantic annotator” to bridge naturalistic movie stimuli, brain responses, and cognitive performance. Using the Human Connectome Project movie-watching dataset, I segmented the film into 293 overlapping clips, prompted Gemini to rate each clip on 11 psychologically interpretable dimensions, and simultaneously extracted clip-wise BOLD activation patterns from the fMRI images in 360 cortical parcels. In this way, the AI and the brain effectively “watch” the same movies in parallel. For each parcel, I then fit linear regression models to predict clip-to-clip variation in movie-evoked responses from these features. Gemini-derived features robustly predicted movie-evoked responses in temporal, medial parietal, and lateral frontal association cortex, but explained little variance in unimodal somatosensory, dorsal parietal, insular, and piriform regions. Feature-weight maps recapitulated known functional specializations, and features with the largest global influence overlapped with the most explainable parcels. Partial least squares analysis revealed that individual differences in resting-state connectivity strength and semantic explainability covaried along an asymmetric intrinsic axis: strongly integrated sensory–opercular systems at rest were associated with poorer AI predictability, whereas a smaller set of dorsal and medial association regions showed enhanced alignment. Finally, regional AI explainability in medial parietal and left perisylvian association areas was positively related to fluid and crystallized cognitive abilities. Together, these findings demonstrate that prompt-defined, interpretable features from foundation models provide a simple and scalable framework for quantifying brain–AI alignment in naturalistic settings, offering a practical bridge between biological and artificial semantic representations.

## INTRODUCTION

Over the past decades, cognitive neuroscience has largely relied on highly controlled, reductionist paradigms using simplified stimuli such as checkerboards, isolated words, or static images to probe the functional architecture of the human brain (1–4). While these designs offer excellent experimental control, they capture only a narrow slice of how the brain operates in everyday life. Over the past decade, however, the field has increasingly turned toward naturalistic paradigms, such as movie watching and narrative understanding, to study the brain in conditions that are closer to real-world experience, where multiple cognitive systems operate together rather than in isolation (5–7). Naturalistic movies and stories elicit robust, distributed, and often highly synchronized BOLD responses across individuals, providing rich opportunities to study large-scale functional organization and intersubject coupling (8,9). However, the same richness that makes movies appealing also creates a methodological bottleneck: unlike traditional event-related designs, movie stimuli present a continuous, high-dimensional stream of visual, auditory, linguistic, and social information that is difficult to translate into a tractable design matrix (5,10,11). As a result, linking the rich neural dynamics observed during naturalistic viewing to the content that participants experience remains a major methodological challenge.

Early movie-fMRI work approached this challenge from the sensory end of the spectrum by extracting low-level visual and auditory features, such as luminance statistics, Gabor or wavelet filter responses, and optical-flow-based motion energy, to quantify the stimulus (12,13). These engineered features successfully explained responses in early sensory cortex, but their predictive power dropped sharply in higher-order regions. This limitation is often referred to as the semantic gap (14–16). As a consequence, much of the neural activity elicited during naturalistic viewing remains unexplained when using feature spaces grounded only in low-level perceptual information. For example, standard optical-flow metrics can capture that “something moves quickly across the screen” but cannot distinguish whether the motion reflects an angry argument or a joyful dance (12,17,18). Thus, purely low-level feature sets provide an important baseline but are intrinsically limited in their ability to account for the semantic and social dimensions that are central to naturalistic experience.

To go beyond purely low-level sensory descriptions of the stimulus, some large-scale naturalistic neuroimaging efforts have adopted ontology-based semantic labeling frameworks, such as semantic category labels derived from WordNet, a hierarchical lexical database widely used in early computer vision and natural language processing research (19,20). Compared with motion-energy and related sensory features, these WordNet-based annotations represent a clear step forward: they provide an explicit, structured vocabulary of objects and actions in the scene (e.g., person, car, walking, speaking. This framework enables more direct comparisons between stimulus content and neural responses in higher-order cortex and has been used to map semantic selectivity during naturalistic movie viewing (21). However, despite this progress, WordNet-based annotation is primarily object- and action-centric. It describes what is visibly present but provides only limited access to the richer psychological dimensions that shape human interpretation, such as conversational context, social tension, narrative structure, or implied mental states. As a result, WordNet still captures only a subset of the constructs that are likely to drive higher-order cortical responses during movie viewing, leaving many layers of meaning unrepresented (21,22).

A third line of work leverages deep neural networks trained on visual or audiovisual tasks, using activations from intermediate layers as high-dimensional embeddings of movie content for encoding or decoding models (23–26). These dense embeddings often capture far more complex structure than hand-crafted low-level features, improving prediction in higher-order cortex by representing objects, scenes, and other abstract attributes beyond pixels and motion vectors. Multimodal deep learning further extends this approach by integrating visual and auditory information to derive comprehensive neural representations of naturalistic stimuli (27–31).

Nevertheless, these representations are typically high-dimensional and opaque: although they can predict brain activity well, they do so in the form of large numerical vectors, often containing hundreds or thousands of values, without clear meaning attached to each dimension. For example, the model may indicate that “dimension 482” or “dimension 917” strongly predicts activity in a specific brain region, yet it remains unclear whether such dimensions represent social interaction, emotional tension, narrative structure, or other latent semantic factors. Current feature-modeling strategies for naturalistic stimuli thus tend to trade off between interpretability (manual, low-dimensional labels) and completeness (high-dimensional, black-box embeddings), leaving a need for representations that are both rich and human-interpretable.

Recent advances in multimodal large language models (LMMs) such as Gemini offer a fundamentally new way to address this problem (32). Trained on massive corpora of text, images, and videos, these models develop integrated world models that can infer objects, actions, social relations, emotions, and narrative context directly from raw visual and auditory input. Given a short movie clip, an LMM can generate detailed, context-sensitive descriptions that resemble human narrative summaries (e.g., “an awkward conversation between coworkers,” “a tense confrontation in a dark hallway”) rather than mere lists of detected objects or undefined numerical vectors. This has led to the hypothesis that LMMs can serve as computational microscopes or “automatic annotators,” transforming qualitative, hard-to-label aspects of naturalistic stimuli into quantitative feature vectors suitable for regression and encoding. When humans watch a movie in the scanner and an LMM “watches” the same movie outside the scanner, the film simultaneously elicits distributed BOLD activity and structured activation patterns within the model. If features derived from the model can be mapped onto the functional topography of the brain, this supports a degree of representational alignment between artificial and biological systems.

Building on these developments, the present study adopts a white-box, explicit scoring strategy for movie-fMRI. Using the Gemini model (https://gemini.google.com) as a multimodal rater, I define a set of eleven psychologically meaningful dimensions, for example, presence of people, degree of social interaction, threat or conflict, and prompt the model to score each short movie segment along these axes. This yields a human-interpretable feature space that captures the high-level semantic and social content while remaining computationally tractable. In parallel, I extract clip-wise fMRI responses from 360 cortical regions of interest in the Human Connectome Project (HCP) 7T movie dataset (33) and fit ROI-wise encoding models that use the Gemini-derived features to predict BOLD activity over time. The resulting regression coefficients and prediction accuracies (R²) form a semantic map of the movie-watching brain, revealing which regions are most sensitive to each feature and how much variance in their activity can be explained by this AI-derived representation. Because every feature has a clear interpretive label, I can move beyond asking what aspects of the movie each brain region encodes.

While encoding models are commonly evaluated at the group level, I hypothesize that the extent to which an individual brain aligns with an AI-derived semantic representation may itself be informative. Large multimodal language models such as Gemini are trained on vast corpora of text, images, and videos, and in doing so, they internalize statistical regularities that reflect shared cultural knowledge and broadly experienced patterns of perception, language, and social meaning. In this sense, such models can be viewed as approximating a population-level representational “center”, a distilled abstraction of how information is typically structured and interpreted across many human experiences and learning histories. If so, the degree to which the movie-evoked brain responses of a subject can be predicted from Gemini semantic features provides a measurable index of how closely their neural representations correspond to this inferred normative space. This perspective raises testable questions: Do individuals whose neural responses are more aligned with the model also demonstrate stronger cognitive performance or more efficient intrinsic network organization? To examine this, for each individual, I relate brain–AI alignment accuracy (R²) to two independent indices of individual variation: intrinsic functional connectivity (FC) measured at rest, and behavioral performance on higher-order cognitive tasks. I found that variability in semantic explainability covaried with resting-state network strength across the cortex. Furthermore, leveraging the cognitive assessments available in the HCP dataset, I observed that individuals whose movie-evoked activity was better predicted by the semantic model tended to have higher fluid intelligence and related abilities.

Taken together, this work demonstrates a principled framework for using multimodal large language models to generate interpretable semantic representations of complex naturalistic stimuli and link them to large-scale neural activity and individual cognitive variation. By pairing explicit LLM-derived semantic dimensions with movie-evoked fMRI responses, I show that artificial semantic structure can be mapped onto cortical organization in a transparent and behaviorally meaningful manner. Moreover, the degree to which an individual brain aligns with this AI-based representation reflects both intrinsic network architecture and cognitive ability, suggesting that alignment may serve as a quantitative marker of representational efficiency. Beyond the specific implementation used here, the framework is extensible: future studies may tailor the semantic space to target constructs such as moral reasoning, theory of mind, episodic memory, or affective dynamics, and employ alternative LLMs or prompting strategies to derive customized representations. In this sense, the approach establishes a two-way bridge, using AI to decode the semantic structure of neural responses under naturalistic experience, and using brain data to test, calibrate, and refine the biological plausibility of advanced AI models. Together, these contributions outline a scalable and interpretable pathway toward studying how meaning is represented in both artificial and biological systems, and how those representational spaces converge or diverge across individuals.

## METHODS

### Participants, Imaging, and Preprocessing

No new MRI data were acquired for this research. Instead, imaging data were downloaded from the Human Connectome Project Young Adult (HCP-Y) database (34). I include all subjects who completed 7T fMRI scans under the movie watching task (n = 176), comprising 70 males and 106 females between the ages of 22 and 35 years.

The proposed analyses will incorporate structural MRI, resting-state fMRI, and task fMRI (movie watching). The imaging protocol has been described in detail elsewhere (35). Briefly, 7T fMRI data were acquired using gradient-echo EPI sequences. Each session consisted of scans with opposing phase encoding directions, with parameters TR = 1000 ms, TE = 22.2 ms, and 1.6 mm isotropic voxel resolution. Importantly, 7T fMRI data are chosen over 3T due to their superior spatial resolution, which is critical for isolating BOLD signals across cortical depth. T1-weighted structural images (3T) were acquired using a 3D MPRAGE sequence (TR = 2400 ms, TE = 2.14 ms, 0.7 mm isotropic resolution). A rich set of behavioral measures is available for these participants and was included in the present analyses. These measures encompass fluid intelligence (PMAT24), episodic memory (PicSeq), working memory (ListSort), cognitive flexibility (CardSort), inhibitory control (Flanker), and composite indices of fluid (CogFluidComp) and crystallized cognition (CogCrystalComp). All scores have been corrected for ages.

T1-weighted images are nonlinearly registered to MNI space using FNIRT (36), and cortical surface reconstructions are generated using FreeSurfer (37). These surfaces were then registered to a standard template using MSMAll (38), which aligns cortical areas based on multimodal features rather than geometry alone. Functional preprocessing includes motion correction, distortion correction (39), and registration to the corresponding structural image, followed by nonlinear alignment to MNI space. After spatial normalization, the data were projected to the standard 32k fs_LR surface mesh and saved in CIFTI dtseries. Additional preprocessing includes denoising via ICA-FIX (40), which removes structured noise components while preserving the neural signal, nuisance regression of motion parameters, linear detrending, and band-pass filtering (0.01–0.1 Hz).

After preprocessing, regional time series were extracted using the HCP-MMP1.0 multimodal parcellation (41), which defines 360 cortical parcels (180 per hemisphere) aligned to the same 32k fs_LR surface space as the functional data. The atlas was applied directly to the CIFTI dtseries files using HCP workbench software. For each participant and each run, this procedure yielded a 360 × T matrix of parcel-averaged BOLD time series, where T corresponds to the number of volumes in the run. Both resting-state and task-derived regional time series were processed using the same pipeline to ensure consistency across analyses. The resulting regional timeseries served as the basis for computing functional connectivity, semantic encoding models, and subsequent individual-differences analyses.

### Movie Information and Clips Generation

Participants completed four movie-watching fMRI runs as part of the HCP 7T protocol: two runs from the CC movie set and two from the HO set. The CC movies consist of short clips drawn from freely available independent films distributed under Creative Commons licensing, whereas the HO movies are composed of professionally edited excerpts from Hollywood films curated and published by a previous study (42). Each run was structured as a compilation of short audiovisual clips that varied widely in emotional tone, narrative structure, visual complexity, and degree of social interaction, providing broad semantic diversity suitable for naturalistic cognitive engagement. Clips were presented sequentially, but each was separated by a 20-second fixation period. These rest intervals temporally isolated consecutive narratives and allowed the hemodynamic response to partially return toward baseline before the next segment began. Participants were given no task instructions other than to watch the movie naturally, and no behavioral responses were recorded. Precise timing information for each clip, including identity, start time, end time, and duration, was obtained from the official HCP metadata and used to align stimulus content to fMRI acquisition timing.

To support time-resolved semantic annotation and encoding analyses, the continuous movie runs were segmented into shorter overlapping clips. Clip generation was performed using a custom MATLAB script and FFmpeg-based video extraction (https://www.ffmpeg.org). For each of the four movie runs, non-rest intervals were first identified based on the official HCP timing tables. Each continuous movie segment was then partitioned into shorter clips using a sliding-window approach (Fig. 1). Clips were targeted to be 20 s in duration, with a minimum allowable length of 16 s to accommodate segments near run boundaries. Consecutive clips were generated with 50% temporal overlap (i.e., 10 s step size), providing dense temporal coverage of the narrative while preserving continuity across neighboring clips. To account for hemodynamic latency and ensure that the extracted visual content corresponded to BOLD time points used in encoding, a 5-second buffer was reserved at the end of each usable segment. All clips were extracted using FFmpeg, recompressed at a standardized frame rate (24 fps), and saved with synchronized audio to preserve semantic context. The final stimulus set contained 293 clips, each associated with metadata including movie run identity, start time, end time, duration, overlap characteristics, and file name. A manifest table (see supporting file for details) documenting all clips and timing information was generated to enable reproducible alignment between video content and fMRI time series.

**Figure 1.**
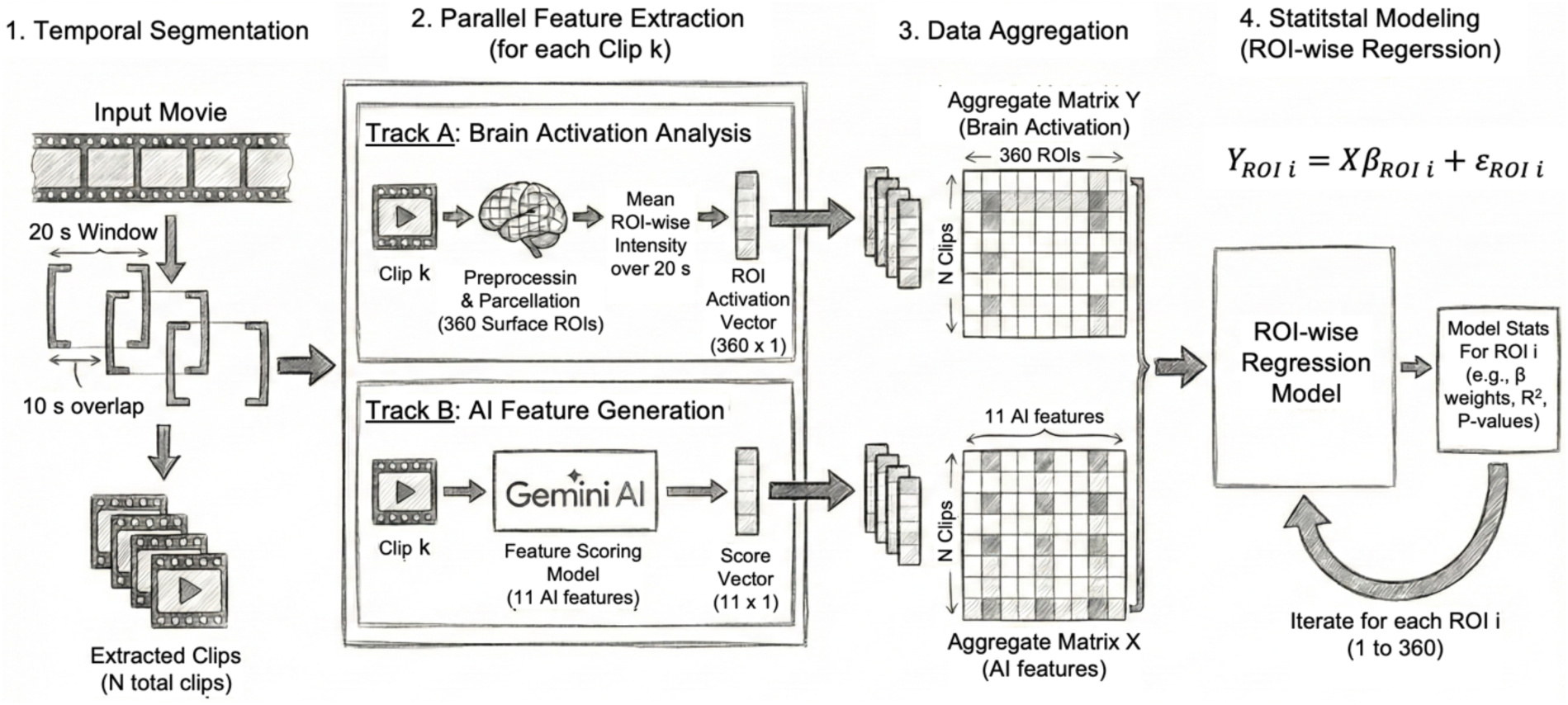
Workflow overview for linking movie-evoked brain activation with AI-derived semantic features. (1) Temporal segmentation. The continuous movie stimulus is segmented into partially overlapping 20-second clips (10-second overlap), producing N clips for analysis. (2) Parallel feature extraction for each clip. Track A: fMRI segment corresponding to each clip undergoes preprocessing and surface-based parcellation into 360 cortical ROIs (HCP-MMP atlas). The mean BOLD intensity over the 20-second window is extracted, yielding a 360×1 activation vector. Track B: Each clip is fed into a multimodal Gemini-based model, which outputs an 11-dimensional semantic feature vector describing visual and narrative properties (e.g., people presence, scene brightness, valence, etc.). (3) Data aggregation. Across all clips, ROI activation vectors form the matrix Y (N clips × 360 ROIs), and AI-derived feature vectors form matrix X (N clips × 11 features). (4) Statistical modeling: ROI-wise regression. For each ROI (i = 1 … 360), a regression model is fit to quantify how well the 11 AI features explain clip-to-clip variability of activation in each ROI. The model yields regression weights (β), goodness-of-fit (R²), and statistical significance for each ROI.

### Clip-wise Brain Patterns

For each participant, preprocessed fMRI data from the four movie runs were available as ROI-wise time series (M, 360 × T), where rows correspond to MMP parcels and columns to TRs. Using the same run-specific timing and rest definitions as in the stimulus design, I first identified non-rest frames by excluding all TRs belonging to 20-s fixation intervals within each run. For each run and participant, I then standardized the ROI time series using z-scoring based on non-rest frames only. To reduce global fluctuations, I removed the frame-wise global mean by subtracting the average signal across ROIs at each TR. For each clip defined in the manifest table (293 clips in total), the corresponding time window was shifted forward by 5 seconds to account for the hemodynamic delay between stimulus onset and BOLD response. For a given clip and participant, I extracted the ROI time series within this lagged window and averaged across TRs, yielding a single 360 × 1 vector representing the mean response pattern for that clip in that individual. To obtain a group-level brain pattern for each clip, I aggregated these participant-specific vectors across all valid subjects. To increase robustness to outliers, I used a 5% trimmed mean across subjects for each ROI (i.e., trimming 5% of the highest and lowest values before averaging). The result is a 360-dimensional group-level response pattern for each of the 293 clips. Importantly, both the group-level clip response patterns and the full set of participant-level clip patterns were stored for downstream analyses, enabling individual-difference analyses.

### Clip-wise AI Features

Semantic features were extracted from each of the 293 movie clips using a custom automated annotation pipeline built around the Gemini multimodal large language model (Gemini 2.5 Pro, Google DeepMind). A lightweight web-based tool was developed in Google AI Studio to streamline clip processing. Video files were converted to Base64-encoded inline data and submitted programmatically to the Gemini API, which returned structured numerical feature scores. The model was chosen for its native ability to process continuous audiovisual input rather than isolated still frames, allowing it to jointly analyze motion, scene context (captions), facial expressions, dialogue, auditory cues, and temporal dynamics. A zero-shot prompting strategy was used to standardize annotations across clips. The instruction asked the model to perform a holistic analysis of each clip and assign a continuous score ranging from 0.0 to 1.0 on eleven predefined dimensions. The feature set was designed to span a gradient from low-level perceptual properties (e.g., motion intensity, scene brightness) to mid-level social and structural cues (e.g., presence of people, dialogue, close-up faces), and finally to higher-order affective and narrative constructs (e.g., valence, arousal, threat, and narrative progress). Clips were processed sequentially to comply with API rate limits, using a queued submission system with a fixed 10-second delay between requests. The final dataset consisted of one semantic feature vector per clip, exported as a structured CSV file for alignment with brain-pattern data and subsequent encoding analyses.

### ROI-wise Semantic Encoding Using AI Features

To quantify the extent to which AI-derived semantic representations predicted neural responses during movie viewing, I constructed ROI-wise encoding models using the clip-level brain patterns and corresponding Gemini-based semantic scores. For each clip, the previously generated group-level, 360-dimensional brain response vector was paired with its semantic feature vector derived from Gemini scoring. All semantic feature dimensions were standardized (z-scored) across clips prior to model fitting. As shown in Fig. 1, Encoding analyses were performed using ridge regression, implemented separately for each of the 360 cortical parcels. For each ROI, I used leave-one-out cross-validation (LOOCV), such that in each fold the model was trained on N–1 clips and tested on the held-out clip. To ensure proper cross-validation hygiene, standardization of both predictors and response variables was performed within each training fold and then applied to the test sample only. The ridge regularization parameter was fixed at λ = 1, chosen to balance model stability and interpretability. Prediction performance was measured as the coefficient of determination (R²) between observed and predicted activity across clips. This procedure produced an R² value for each of the 360 ROIs, yielding a spatial map of semantic explainability across the cortex. In addition to prediction accuracy, I quantified feature contribution by examining the magnitude of regression coefficients. For each feature, I computed the mean absolute coefficient across ROIs as a measure of overall influence, and a second weighted metric in which coefficients were weighted by the ROI-specific R² values, emphasizing features that contributed most to regions with high model predictability.

### Resting-State FC Strength and Its Association With AI–Brain Encoding Performance

To examine whether intrinsic network organization relates to how well an individual brain aligns with AI-derived semantic features, I quantified resting-state FC strength for each subject and assessed its association with regional encoding performance (R² values from movie-based ridge encoding). Resting-state fMRI data for each participant were processed using the same standardized pipeline applied to the movie runs. Time series were extracted from 360 cortical regions of interest (ROIs) defined by the MMP parcellation. For each subject, I computed a 360 × 360 FC matrix by estimating pairwise Pearson correlations across ROI time series. The resulting matrices were Fisher-z transformed to stabilize variance. To derive a subject-level measure of intrinsic network organization, I computed a nodewise connectivity strength metric for each region, defined as the mean Fisher-z correlation (negative FCs were set to 0) between that region and all other ROIs. This resulted in a 360-element vector per participant, capturing how strongly each cortical region is functionally integrated with the rest of the cortex during rest.

I next examined whether the spatial pattern of intrinsic connectivity strength relates to how explainable the activity of each region is by the Gemini-derived semantic features. I employed Partial Least Squares (PLS), a multivariate technique, to statistically assess the correspondence between encoding performance and FC strength. In this analysis, subjects formed the rows of the data matrix (N subjects), and ROIs were the shared variable dimension (360 features). The two input matrices were:

X: ROI-wise resting-state FC strength (subjects × 360)

Y: ROI-wise R² encoding performance (subjects × 360)

PLS identifies latent components that maximize the covariance between X and Y. The first latent variable (LV1) was interpreted as the dominant axis linking intrinsic network organization to AI explainability. The statistical significance of LV1 was assessed using permutation testing (1,000 permutations), and the reliability of ROI loadings was evaluated with bootstrap resampling, yielding z-scored stability maps. Regions with stable positive weights on LV1 were interpreted as areas where stronger intrinsic connectivity was systematically associated with higher AI–brain alignment.

### ROI-wise Association Between AI Encoding Performance and Cognitive Scores

To test whether brain–AI alignment relates to individual cognitive abilities, I correlated ROI-wise encoding performance with behavioral measures across subjects. For each participant, I obtained a 360 × 1 vector of encoding performance (R²) from the ridge regression models described above. These values quantify, for each MMP parcel, how well the movie-evoked responses of that subject are predicted by the Gemini-derived semantic features. For behavior measures, I focused on a set of canonical HCP cognitive indices (age-adjusted), including fluid intelligence (PMAT24), episodic memory (PicSeq), working memory (ListSort), cognitive flexibility (CardSort), inhibitory control (Flanker), age-adjusted fluid cognition composite (CogFluidComp), and crystalized cognition composite (CogCrystalComp). I constructed a 360 × N matrix of encoding performance (R²) and an N × 7 matrix of cognitive scores, where 7 is the number of available cognitive measures. For each ROI and each cognitive variable, I computed a cross-subject Spearman rank correlation between the R² values and the cognitive scores. This produced a 360 × 11 matrix of correlation coefficients (r) and corresponding raw p-values (p). To control for multiple comparisons across all ROI × cognition pairs, I applied a Benjamini–Hochberg false discovery rate (FDR) procedure at q = 0.05, using all correlation p-values pooled into a single vector, and significant effects were defined as FDR-corrected p < 0.05.

## RESULTS

### Regional Variability in Semantic Predictability Across the Cortex

To determine how well high-level semantic information extracted from the movie clips accounted for regional brain responses, ROI-wise ridge regression models were fit for all 360 cortical parcels in the MMP atlas. Cross-validated prediction performance was quantified using the coefficient of determination (R²), reflecting how much clip-to-clip variance in movie-evoked activity could be explained by the 11 AI-derived semantic features. As shown in Figure 2a, prediction accuracy varied widely across the cortex, revealing distinct spatial patterns rather than uniform explanatory power. The highest R² values emerged in a distributed set of association regions, including superior temporal sulcus (STS), middle and anterior temporal cortex, precuneus, posterior cingulate cortex, and lateral prefrontal association areas. The distribution of R² values across ROIs (Figure 2b) further highlights strong regional heterogeneity. While the majority of parcels exhibited modest but positive prediction values, a subset reached relatively high explanatory performance, exceeding 0.30. A small number of parcels yielded values near zero or slightly negative, the latter reflecting cases where the cross-validated model performed marginally worse than a mean-only baseline. Ranking the parcels confirmed these trends. The top-10 most predictable regions (L and R STGa, L and R A5, L STSdp, STSda and STSva, L TGd and TGv, and L 55b; Fig. 2c) were located almost entirely in the superior temporal gyrus/sulcus and anterior temporal association cortex, with an additional dorsal frontal premotor parcel (55b) in lateral frontal cortex. Together, these regions belong to networks implicated in higher-order auditory processing, speech and language, and social–narrative understanding. By contrast, the bottom-10 ROIs (R and L 5L, L i6-8, R 7PC, L Pol2, R 2, L OP1, L IP1, L 7Pm and L Pir; Fig. 2d) included primary and secondary somatosensory areas (2, 5L, OP1), dorsal and inferior parietal association regions (7PC, 7Pm, IP1), a dorsolateral frontal control parcel (i6-8), an posterior insular area (PoI2), and piriform cortex (Pir). These parcels are mainly associated with somatosensory, sensorimotor, attentional/control, visual, or olfactory processing, and therefore show little variance explained by the high-level semantic features modeled here.

**Figure 2.**
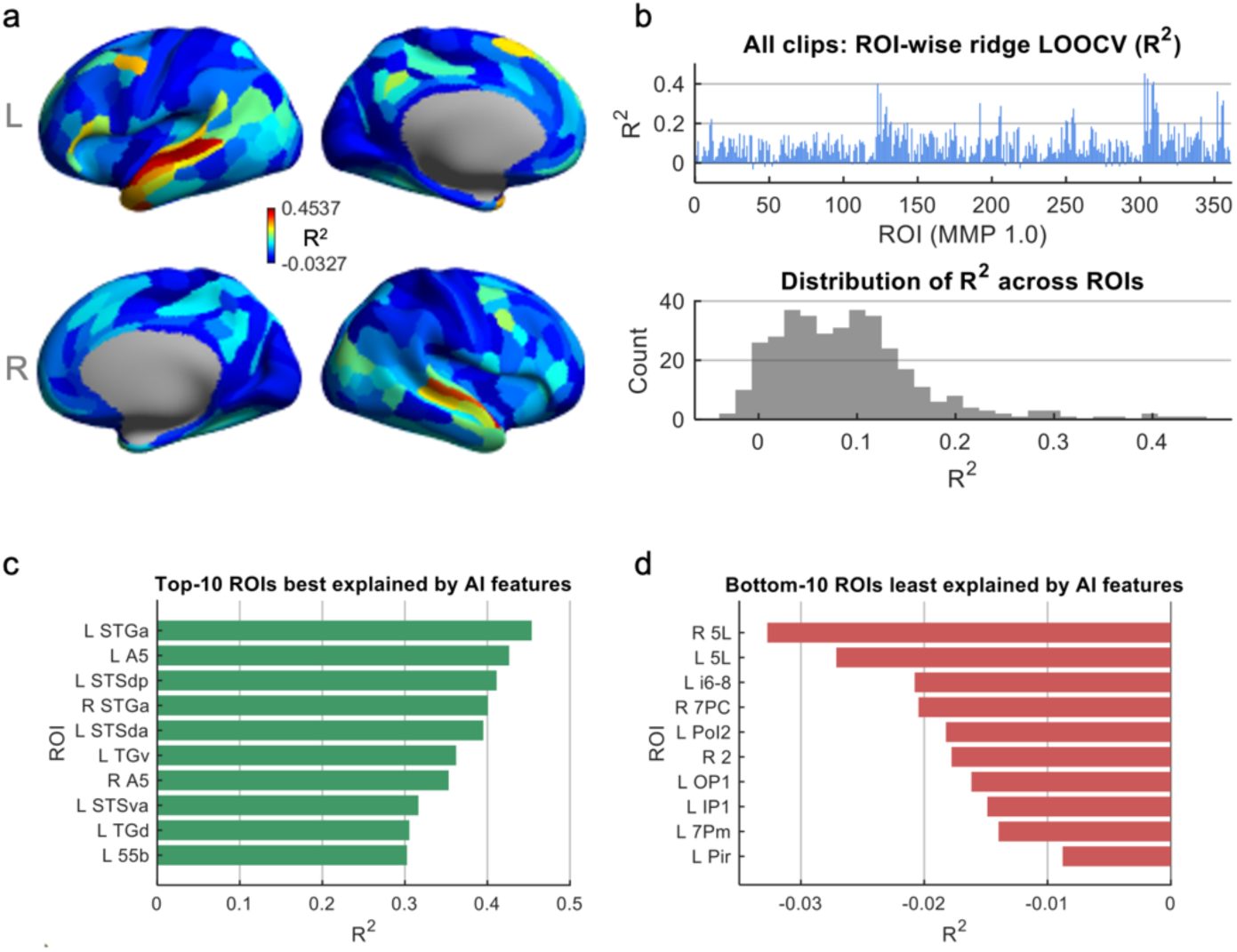
ROI-wise predictive performance of AI features across the cortex. (a) Surface maps of prediction accuracy (group mean). Each parcel is colored by its cross-validated coefficient of determination (R²) from the ROI-wise ridge regression model, reflecting how well the 11 AI-derived movie features explain the 20-s windowed brain activation patterns. Warmer colors indicate ROIs whose activity patterns are more strongly predicted by the feature set. (b) Distribution of predictive performance across all ROIs. Top: ROI-wise R² values for all 360 MMP parcels. Bottom: Histogram summarizing the distribution of R² across the cortex, showing substantial regional variability. (c) Top-10 most predictable ROIs. Bar plot showing the 10 parcels with the highest cross-validated R² values, indicating regions whose movie-evoked activity is best captured by the AI feature set. (d) Bottom-10 least predictable ROIs. ROIs with the lowest (or slightly negative) R² values, representing regions whose activity patterns are poorly explained by the available AI features.

### Contribution of Semantic Feature to The Prediction Model

To characterize how different semantic dimensions from the Gemini model are represented across the cortex, ridge-encoding coefficients were converted to standardized absolute weights (|β|) for each feature and each of the MMP parcels. The surface maps in Figure 3a–b visualize these |β| values, with higher values indicating stronger sensitivity of the movie-evoked pattern of a parcel to a given feature. The People Presence feature showed its strongest weights in bilateral TPOJ3 (temporo-parieto-occipital junction, multimodal association cortex), TF (ventral temporal fusiform cortex), and LO3 (lateral occipital visual association cortex). These areas lie at the interface of occipital, temporal, and parietal regions and are commonly implicated in high-level visual analysis of people and objects, consistent with a sensitivity to the presence of humans in complex scenes. The Faces Close-Up feature exhibited pronounced weights in right area 44 (inferior frontal cortex) and dorsal/posterior visual regions, including V6A, VMV1, and VMV2. These areas span frontal regions involved in orofacial and speech-related motor representations and medial/ventral occipital regions associated with detailed visual analysis. Social Interaction weights concentrated in left 31a (posterior cingulate/retrosplenial default-mode area) and parahippocampal regions PHA2 and PHA3, with corresponding peaks in right 31a and PHA3. This pattern aligns with a medial temporal–posterior cingulate network involved in contextual, episodic, and social–scene processing. The Dialogue feature showed the largest single coefficients across the entire matrix in left A5 and bilateral STGa, complemented by high weights in left STSdp and right A5. These regions are located in the superior temporal gyrus and sulcus and adjacent belt areas and are classically associated with higher-order auditory and speech processing. For Music, the strongest coefficients appeared in bilateral TA2 (auditory association cortex), left 31a (posterior cingulate), left POS1 (parieto-occipital sulcus), and right s32 (ventromedial prefrontal cortex), consistent with a distributed network spanning auditory, medial parietal, and medial prefrontal regions engaged by musical and affective context. Several features recruited more frontally distributed systems. Motion Intensity showed its largest weights in bilateral MI (middle insular cortex) and frontal opercular regions FOP4, together with the dorsal medial frontal area p32pr. This pattern corresponds to cingulo-opercular and insular control networks often linked to salience and integrated sensory–motor processing, indicating that perceived motion intensity in the clips is encoded in multimodal control and interoceptive hubs rather than early visual cortex alone. Scene Brightness weights were concentrated in left medial frontal and parietal midline regions, including 9p, 7m, 6mp, 31pv, and 31pd, which overlap with default-mode territories along medial prefrontal and posterior cingulate cortex. Affective and appraisal-related features showed distinct prefrontal– limbic signatures. Valence assigned high weights to right 9-46d and 46 (dorsolateral prefrontal regions), left 47m (orbitofrontal cortex), left PeEc (perirhinal/ectorhinal cortex in medial temporal lobe), and right 23c (posterior cingulate). This combination of dorsolateral, orbitofrontal, and medial temporal areas is consistent with networks involved in evaluating the emotional meaning and value of complex events. Arousal weights were maximal in right 10v and 10r (ventral anterior prefrontal areas) and right 8Ad, together with left 10r and right STSva, indicating a ventromedial and dorsal frontal network, plus superior temporal association cortex. Threat weights localized primarily to left orbitofrontal and lateral prefrontal parcels a47r, p47r, 10pp, 8Av, and 8BL, regions associated with evaluation of potential risk, punishment, and behavioral control. Finally, Narrative Progress was most strongly expressed in cingulo-opercular and dorsal frontal eye–field regions, including SCEF, 24dd, bilateral 6mp, and FEF, consistent with a network supporting sustained control, monitoring, and alignment of attention as the story advances.

**Figure 3.**
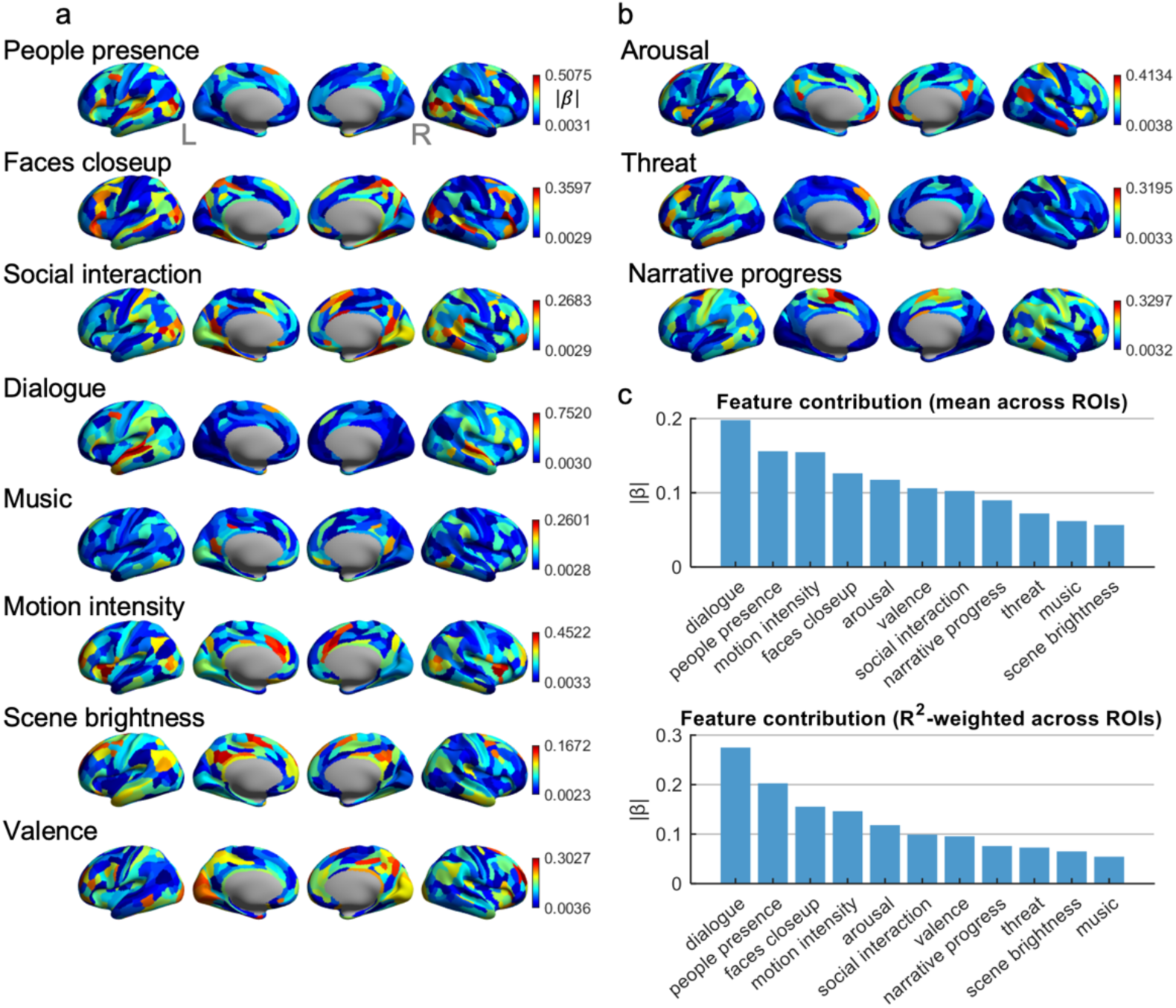
ROI-wise regression weights for individual AI features and their overall contributions. (a) and (b) Cortical maps of regression coefficients (|β|) for eleven semantic–perceptual AI features. Results show the brain-wide spatial distribution of standardized regression weights (absolute value of β) for each feature when predicting ROI-wise activation across all movie clips. Warmer colors indicate stronger associations between a given AI feature and the activation pattern of each cortical region. (c) Feature-level contributions across all ROIs. Top: Mean |β| across 360 ROIs, reflecting the overall influence of each feature on explaining brain activation patterns. Bottom: R²-weighted |β|, which up-weights features that explain ROIs with higher cross-validated predictive accuracy, providing an importance score aligned with model fit.

Panel 3c summarizes the overall contribution of each feature by averaging |β| across all ROIs (and weighting by ROI-wise prediction performance). Dialogue showed the largest overall contribution to explaining cortical patterns, followed by People Presence and Motion Intensity, with Faces Close-Up, Arousal, and Valence contributing at intermediate levels and Music, Threat, Narrative Progress, and Scene Brightness contributing more modestly. Together, these results indicate that the AI-derived semantic space is not uniformly expressed across the cortex: features tied to speech, social agents, and dynamic action account for a substantial portion of explainable variance, whereas features such as brightness or low-level soundtrack properties play a comparatively smaller role in shaping the distributed movie-evoked patterns in this dataset.

### Brain–AI Alignment Axis Linking Semantic Explainability to Intrinsic Connectivity

To relate individual differences in AI explainability to intrinsic functional architecture, I applied partial least squares (PLS) to two ROI × subject matrices: (i) semantic explainability (R²) from the encoding models, and (ii) resting-state FC strength for the same MMP parcels. The first latent variable (PLS Axis 1) captured the dominant mode of covariation between these two profiles across participants (Fig. 4). At the subject level, X-scores (R² side) and Y-scores (FC-strength side) were positively correlated (r = 0.52; Fig. 4c), indicating that individuals whose cortical activity patterns were better predicted by the AI-derived semantic features tended to occupy a similar position along an intrinsic connectivity gradient. Axis 1 also explained the largest share of variance in both spaces (Fig. 4b), making it the primary “brain–AI alignment” dimension examined here.

**Figure 4.**
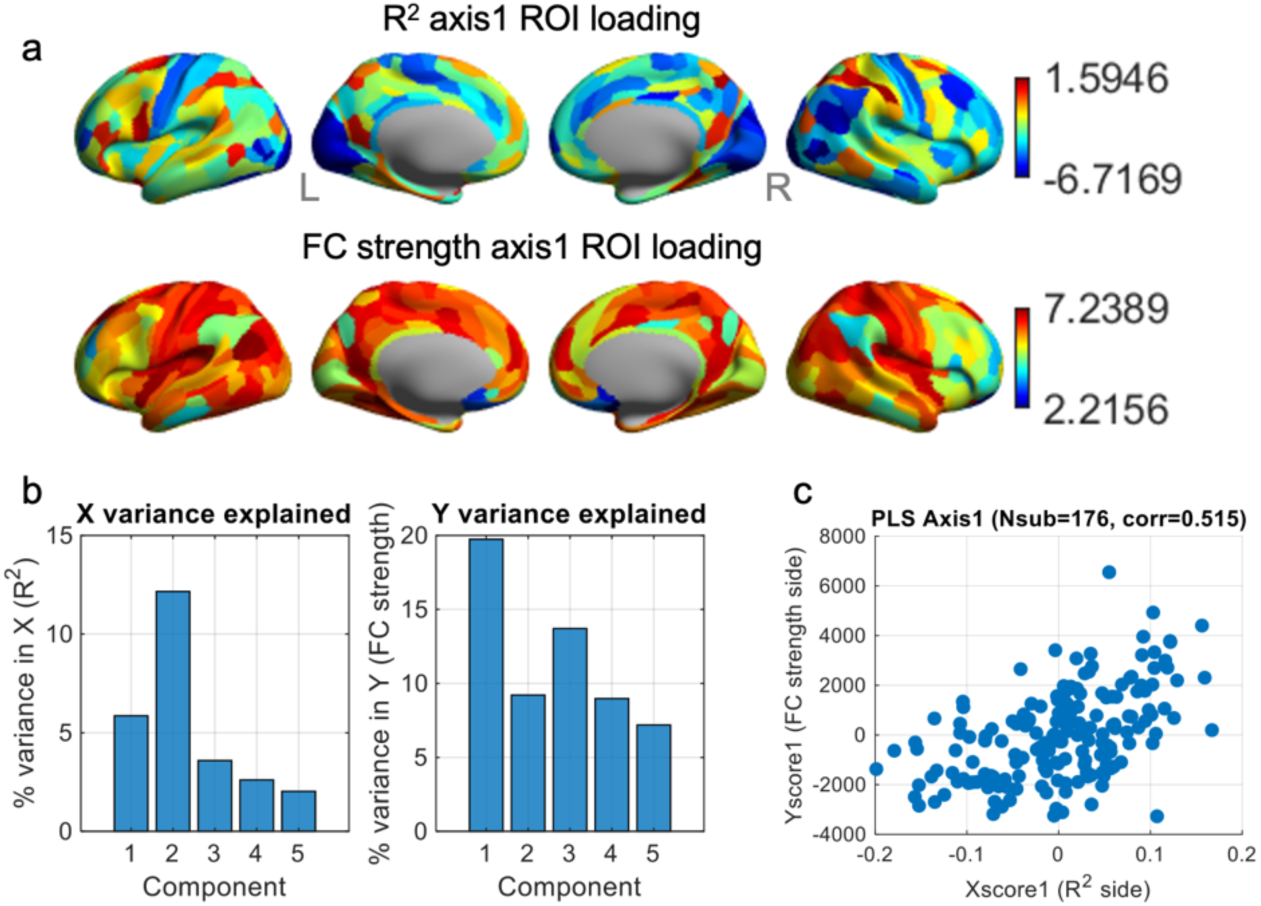
Partial least squares (PLS) analysis linking ROI-wise R² profiles and rsFC-strength profiles across subjects. (a) PLS Axis 1 ROI loadings. Top row: spatial loading pattern for the first latent variable (Axis 1) derived from the across-subject R² profiles (how well AI features explain clip-wise brain activation patterns of each subject). Bottom row: corresponding loading pattern for rsFC-strength profiles. Warm colors indicate ROIs with strongly positive loadings; cool colors indicate strongly negative loadings. These loadings reflect ROIs that contribute most to the covariation between R² and FC-strength patterns across subjects. (b) Variance explained by PLS components. Left: percentage of variance in the R² side (X) explained by each PLS component. Right: percentage of variance in the FC-strength side (Y) explained by each component. Component 1 explains the largest share of variance on both sides, indicating the dominant cross-subject mode of R²–FC covariation. (c) Scatterplot of X-scores (R² side) versus Y-scores (FC-strength side) for all subjects.

The spatial loading patterns for Axis 1 revealed a striking asymmetry between the two modalities (Fig. 4a). On the R² side, loadings were largely shifted toward negative values (range: −6.7 to +1.6, median: -2.8), with relatively few parcels showing strongly positive contributions. The most positive R² loadings were found in right dorsal and lateral association cortex, including medial intraparietal area (MIP), anterior intraparietal area (AIP), and inferior parietal areas PFt and 7PL, as well as posterior cingulate/precuneus parcel POS1. In these regions, subjects with stronger resting FC along Axis 1 also tended to show higher AI explainability, reflecting a modest “aligned” mode where a more strongly connected dorsal attention, parietal–cingulate backbone accompanies better capture of movie-evoked responses by the semantic model. In contrast, many sensory and sensorimotor regions exhibited strongly negative R² loadings while simultaneously showing high positive FC-strength loadings on the same axis. This was particularly evident in early visual and motion-sensitive cortex (e.g., V1, V2, V3, V3A, MT, MST, FFC), dorsal mid-cingulate area 24dv, somatomotor regions (areas 3a, 1, 5m), and insular/opercular parcels (e.g., FOP4, PoI1/2). For these ROIs, individuals with stronger intrinsic connectivity along Axis 1 tended to show lower semantic explainability, yielding a pronounced “anti-aligned” component of the solution. The FC-strength loadings themselves were almost uniformly positive (range: 2.2 to 7.2, median: 6.0), with peaks in medial temporal (L PHA1), inferior parietal (L PGi), dorsal prefrontal (L 8Ad), and posterior insular/opercular regions (L/R PoI2, L OP2-3), as well as ventromedial visual cortex (L VMV2).

### Brain–AI Explainability Predicts Individual Differences in Cognitive Performance

I next asked whether individual differences in brain-AI alignment were related to cognitive performance (Figure 5). For each subject and each ROI, I correlated the movie-encoding R^2^ with seven HCP cognitive scores. After FDR correction (q < 0.05) across all ROI-cognition tests, significant associations emerged in specific, highly localized networks (Figure 5b). Fluid intelligence (PMAT24) was significantly positively associated with explainability in left area 45 (inferior frontal gyrus/Broca’s area) and left STSvp (superior temporal sulcus), while showing a negative association with left FOP1 (frontal operculum). Working memory (ListSort) performance showed strong positive associations with explainability in the posterior cingulate and medial parietal cortex, specifically bilateral area 31pd and left 31pv, as well as left STSvp. For Inhibitory Control (Flanker), I observed a single significant negative association in left area 5L (somatosensory cortex). The Fluid Cognition Composite (CogFluidComp) aligned with the individual fluid measures, showing its only significant peak in left 7Pm (medial parietal cortex). Finally, the Crystallized Cognition Composite (CogCrystalComp) showed the most robust effects, with the strongest positive correlations found in left medial parietal regions L 5mv and L 23c, left STSdp and left PGi, as well as R 5mv. Explore Together, these results reveal a consistent topography: individuals whose neural responses in medial parietal and left-lateralized language/social association areas are better predicted by the AI semantic features tend to demonstrate superior cognitive performance.

**Figure 5.**
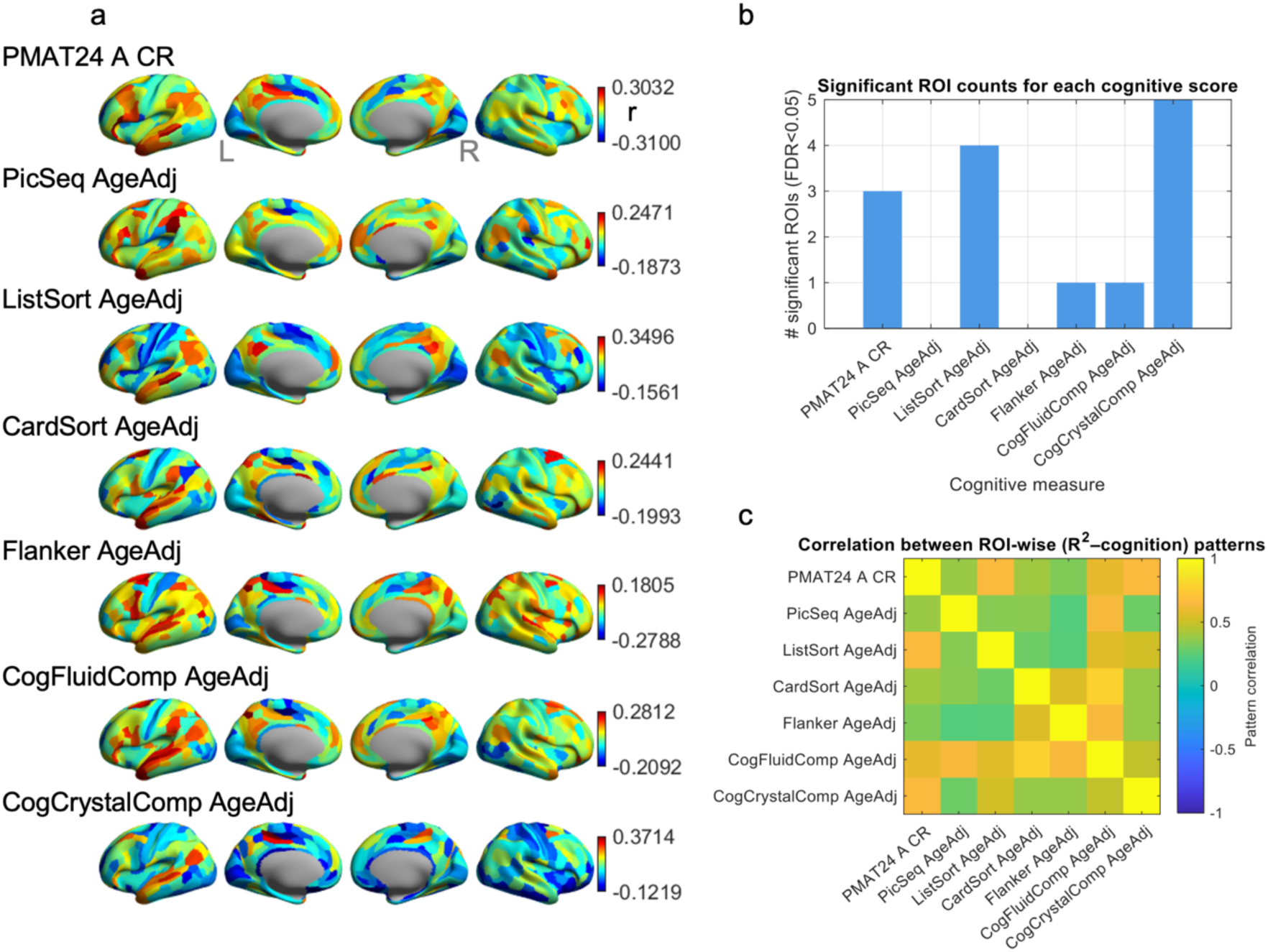
Associations between ROI-wise explainability (R²) and individual cognitive abilities. (a) ROI-wise Spearman correlations between explainability and cognition. Surface maps show the resulting r values for each ROI (MMP 1.0 parcellation). Warm colors denote positive associations (higher cognitive score associated with higher explainability), while cool colors denote negative associations. (b) Number of significant ROIs for each cognitive measure. For each cognitive score, the number of ROIs showing a significant correlation with R² (FDR < 0.05 across all ROI × cognitive tests) is plotted. (c) Similarity between explainability–cognition spatial patterns. A correlation matrix showing pairwise pattern similarity between the ROI-wise correlation maps of all cognitive measures. Each entry reflects the Pearson correlation between two cognitive scores’ r-maps from panel (a).

Despite the sparsity of FDR-significant ROIs, the spatial patterns of brain-AI-cognition coupling were systematically related (Figure 5c). The ROI-wise r maps for fluid measures (PMAT24, ListSort, CogFluidComp) were strongly correlated with each other, sharing a medial-parietal profile. These results suggest that individuals whose cortical activity, particularly in medial parietal, posterior cingulate, and right lateral association regions, is better captured by the Gemini-derived semantic features tend to show stronger performance on higher-order cognitive tests.

## DISCUSSIONS

In this study, I used a multimodal large language model as an automated “semantic annotator” to link naturalistic movie content with large-scale patterns of human brain activity. By having Gemini generate a compact set of perceptual, social–affective, and narrative features for hundreds of overlapping clips from HCP 7T movies, I showed that these AI-derived descriptors robustly explained time-resolved cortical responses across a broad association network, while leaving many primary sensorimotor and insular regions largely unexplained. The feature-weight maps revealed distinct topographies for different dimensions of experience. At the individual level, a multivariate analysis revealed a pronounced inverse relationship in unimodal cortex: subjects whose resting-state networks were more strongly functionally integrated tended to show lower brain–AI semantic alignment in primary sensory, motor, and insular regions, even as alignment remained concentrated in higher-order association cortex. Finally, individual differences in explainability within specific parcels were selectively related to fluid and crystallized cognitive abilities. Together, these findings provide an initial demonstration that LLM-based video annotations can serve as a powerful, scalable bridge between the semantic structure of complex real-world stimuli, intrinsic brain network organization, and variation in cognitive performance across individuals.

### AI–Brain Semantic Alignment Concentrated in the Association Cortex

A first key insight from these analyses is that AI–brain predictability is highly non-uniform across the cortex, and it closely follows known functional hierarchies. The LLM-derived feature space yielded the highest explainability in transmodal association regions along the superior temporal, anterior temporal, medial parietal, and lateral frontal cortices, whereas primary sensory, somatomotor, insular, and piriform areas were only weakly predicted. Consistent with this view, the parcels with the highest R² carried substantial weights for the most influential features, particularly dialogue, people presence, and motion intensity, indicating that naturalistic speech, social content, and dynamic visual events are the main drivers of movie-evoked responses in these networks. Rather than being tuned to a single feature, however, these hubs showed clear mixed selectivity, with coordinated contributions from multiple social, affective, and narrative dimensions, in line with the idea that high-dimensional mixed selectivity is a hallmark of association cortex supporting flexible cognition (43,44). By contrast, the poor predictability in somatosensory, motor, insular, and olfactory cortices likely reflects, first and foremost, the fact that the HCP movie paradigm provides relatively weak and indirect stimulation of these systems, for example, very limited actual bodily sensation, interoceptive change, or olfactory input, so that there is little reliable variance for any semantic model to capture. A second, complementary possibility is that the current 11-dimensional feature set under-represents the specific low-level and bodily codes that these regions carry, which are not explicitly modeled by the LLM prompt. Together, this divergence highlights both the strength and the limitation of using a compact, interpretable AI feature space: it captures the dominant high-level psychological drivers of movie-evoked activity in transmodal networks but likely misses aspects of low-level and bodily processing that may only emerge under richer task manipulations and with more specialized feature sets in future work.

### Intrinsic network architecture shapes AI–brain alignment

The PLS analysis relating resting-state connectivity strength to AI-based semantic explainability shows that individual differences in brain–AI alignment are not only a property of movie-evoked responses but are also rooted in the intrinsic functional architecture of the brain. Resting-state FC is often taken as a trait-like marker of how cortical regions are wired to interact in the absence of explicit task demands (45), and here the first PLS axis captured a dominant “brain–AI alignment” dimension: the degree to which an individual brain approximates the AI-defined semantic space during naturalistic viewing is systematically constrained by its resting-state network organization. Importantly, however, the ROI-level pattern of PLS weights indicates that this axis does not simply reflect a uniform “more connectivity, more explainability” relationship. Axis 1 is dominated by a large set of visual, motion-sensitive, somatosensory, and insular/opercular regions that show strongly positive weights for FC strength but robustly negative weights for AI explainability. In other words, individuals with more strongly unimodal networks at rest tend to show less of their movie-evoked variance in these regions captured by the Gemini-derived features. In contrast, a smaller cluster of dorsal parietal and medial posterior association regions (including superior parietal and posterior cingulate/medial parietal parcels) shows a more classical positive–positive pattern, where greater intrinsic connectivity strength goes along with higher semantic predictability.

Thus, the axis is highly asymmetric, and this opposing pattern between unimodal and association regions closely follows well-established macroscale gradients from sensorimotor to transmodal cortex, where unimodal systems at one end carry concrete sensory and bodily codes, while default-mode and higher association networks at the other end support abstract, integrative representations (46–48). One possible explanation is that Axis 1 largely reflects a redistribution of representational emphasis along that gradient. In individuals whose intrinsic architecture places more “weight” on sensory and opercular systems, movie-evoked responses in those regions may be dominated by low-level signals that fall outside the current 11-dimensional semantic feature set, producing strong FC but weak AI explainability. Conversely, individuals nearer the transmodal end of the axis may express more stable, high-dimensional mixed representations in dorsal and medial association cortex that align more directly with the AI feature space, while unimodal regions play a relatively smaller role in the shared variance captured by the model. Together, these findings highlight that resting-state network architecture does not simply scale up or down the overall strength of AI–brain correspondence but selectively shapes where in the cortical hierarchy semantic predictability emerges. Future work could build on this by decomposing intrinsic connectivity within and between specific networks, examining dynamic connectivity states during naturalistic viewing, and testing whether similar asymmetric patterns arise when semantic features are expanded to better cover low-level and bodily dimensions.

### Brain–AI alignment in the association cortex relates to cognitive ability

The associations between ROI-wise explainability and cognition indicate that brain–AI alignment is not only a descriptive property of movie-evoked responses but also carries behavioral relevance. Although the number of FDR-significant ROIs was modest, the effects were spatially specific and converged on a coherent network profile. Across all measures, individuals whose neural responses in medial parietal and left-lateralized language/social association areas were better predicted by the AI semantic features tended to demonstrate superior performance on higher-order cognitive tests. This topography aligns well with prior work implicating medial parietal/posterior cingulate cortex and lateral temporal–parietal association areas in semantic cognition, memory, and complex reasoning (49–51). The observation that explainability in these regions relates to both fluid and crystallized cognition suggests that what matters for cognitive performance is not simply how strongly they activate, but how efficiently their movie-evoked patterns can be captured by a compact, semantically interpretable feature space. In other words, individuals whose association areas express neural responses that are “well-behaved” with respect to the AI semantic axes tend to perform better on demanding reasoning, working-memory, and vocabulary tasks. These findings dovetail naturally with the “AI as center” perspective I hypothesized. Large multimodal models such as Gemini are trained to approximate a distilled abstraction of how information is typically structured and interpreted across many human experiences (52,53). Therefore, brain–AI alignment, operationalized as R² for movie encoding, indexes how closely neural representations during naturalistic viewing of an individual approach this inferred normative semantic space.

I addition, the pattern is neither global nor uniformly positive. For the Flanker inhibitory-control task, the correlations between explainability and performance were modest overall but skewed toward negative values (roughly –0.27 to 0.18), indicating that better inhibitory control tended to be associated with slightly lower AI-based predictability, especially outside the association hubs highlighted by the other cognitive measures. Within the present framework, this should not be viewed as a “worse” brain state. Instead, it could suggest that the neural patterns supporting strong inhibition are less directly expressed along the compact semantic space defined by the Gemini features and are therefore harder for this particular model to capture. Given the small effect sizes and sparse pattern, we treat this as a tentative observation and place greater emphasis on the more robust positive associations.

Finally, the overall patterns of R²–cognition correlation maps for fluid measures (PMAT24, ListSort, CogFluidComp) are highly similar and share a medial parietal profile, indicating that the effects concentrate in a consistent, theoretically meaningful network. This suggests that brain–AI alignment is most behaviorally relevant when it occurs in specific association hubs, rather than being uniformly high across the cortex. Future work with larger and more diverse samples could test the robustness and generality of these associations, examine longitudinal links between alignment and cognitive change, and extend the framework to other domains by tailoring the AI feature space. In this way, R²–cognition relationships provide not only a behavioral validation of the AI-derived semantic representation, but also a potential quantitative marker of how efficiently key cortical hubs implement a normative, center-like organization of meaning.

### LLM-Based Annotations as a Bridge Between Biological and Artificial Intelligence

Beyond the specific findings, this work illustrates a general strategy for using large multimodal models as a bridge between human intelligence and artificial intelligence. Rather than treating Gemini as a black box that produces high-dimensional, opaque embeddings, I used it to generate a small set of explicit, interpretable features that can be directly related to psychological constructs and cortical topographies. Similar to recent proposals that derive interpretable axes by asking LLMs structured questions or use LLM text embeddings to model high-level visual and semantic representations in the brain (54,55), this approach positions foundation models as flexible “annotation engines” that expose their internal semantic structure in a form that is both neurobiologically meaningful and statistically tractable. In this sense, the AI model provides a population-level representational template, while the brain data reveal where and how individual nervous systems approximate, diverge from, or reorganize around that template.

At the same time, it is important to acknowledge that the present 11-dimensional feature space is necessarily incomplete. Many aspects of movie experience, fine-grained visual form, detailed motor plans, interoceptive and visceral states, and long-range narrative dependencies, are not fully captured by these few semantic axes, and this likely contributes to the low predictability observed in unimodal and insular regions. Moreover, the neural patterns I model are derived from clip-wise BOLD changes using a sliding-window scheme that is closely related to the dynamic-variation approaches often applied to resting-state data (56–58). This representation is well suited to continuous movie stimulation, which, like rest, lacks clear trial boundaries and unfolds over long timescales, but it inevitably compresses the underlying dynamics and may miss faster or more complex BOLD fluctuations that are only partially coupled to the current semantic features. Future work could combine LLM-based annotations with richer models of brain dynamics to better capture the full temporal structure of movie-evoked activity. This limitation is also a strength. Because the features are defined by simple natural-language prompts, the framework is highly adaptable to different study designs. Researchers can easily re-prompt the model to emphasize other constructs (e.g., threat types, social hierarchy, symptoms, or clinical behaviors) or expand the feature set when richer coverage is needed, much like constructing task-specific but interpretable embeddings for different cognitive domains. A practical advantage of this pipeline is that it is technically lightweight and accessible, even for investigators without a software-engineering background. Using Google AI Studio and the Gemini API, semantic scoring of long videos can be implemented with only a few lines of short, human-readable prompts, with the platform handling video ingestion, multimodal processing, and model hosting behind the scenes. This lowers the barrier for clinical and translational researchers who may not have the resources to build bespoke computer-vision models, but who routinely collect rich behavioral, neuroimaging, or symptom data that could benefit from standardized, scalable annotation of complex stimuli.

Looking ahead, this kind of AI–brain interface suggests a two-way relationship rather than a one-directional borrowing of AI models to explain neural data. On the one hand, foundation models offer a powerful toolset for cognitive and clinical neuroscience, providing high-coverage, customizable descriptors of naturalistic stimuli and candidate representational spaces for interpreting brain activity. On the other hand, the brain can serve as a rich source of constraints and training signals for improving AI itself, especially in domains where current models still struggle: long-timescale integration, high-order abstraction, and durable memory (59–61). For example, default-mode and medial temporal networks integrate emotional and narrative information over tens of minutes during movies, linking the affective tone of a scene to events that occurred many scenes earlier. Embedding such brain-inspired temporal and hierarchical principles into future multimodal models could help them move beyond local, short-context descriptions toward a deeper understanding of extended real-world events. In this view, interpretable, LLM-based annotations are not only a convenient tool for studying the brain but also a stepping stone toward a tighter, mutually informative dialogue between biological and artificial intelligence.

### Conclusions

This study shows that a large multimodal language model can be used as an interpretable bridge between naturalistic movie content, brain activity, and cognition. Gemini-derived semantic features robustly predicted movie-evoked responses in temporal and medial parietal association cortex, while largely failing to explain unimodal and insular regions, and individual differences in resting-state network organization selectively shaped where this AI–brain alignment emerged. Explainability in medial parietal and left perisylvian association areas was further related to fluid and crystallized cognitive abilities, indicating that alignment with an AI-derived semantic space in specific hubs is behaviorally meaningful. Although the current 11-dimensional feature set is incomplete, its prompt-based, easily customizable design offers a flexible and accessible framework that future work can extend to richer features, additional brain systems, and tighter, bidirectional links between biological and artificial intelligence.

## DATA AVAILABILITY

The MRI data used in this study are available in the HCP database https://www.humanconnectome.org/

## CODE AVAILABILITY

Software and toolboxes that are used in this study:

Gemini: https://gemini.google.com/

Google AI Studio: https://aistudio.google.com/

FFmpeg: https://www.ffmpeg.org/

CIFTI: https://www.nitrc.org/projects/cifti/

GIFTI: https://www.nitrc.org/projects/gifti/

HCP workbench: https://www.humanconnectome.org/software/connectome-workbench

## ACKNOWLEDGEMENTS

I am grateful to Dr. John Gore for his continuous support, mentorship, and insightful discussions throughout this work. I also thank Dr. Zhaohua Ding for valuable feedback on the analyses and interpretation of the results.

Data of young adults were provided by the Human Connectome Project, WU-Minn Consortium (Principal Investigators: David Van Essen and Kamil Ugurbil; 1U54MH091657), funded by the 16 NIH Institutes and Centers that support the NIH Blueprint for Neuroscience Research; and by the McDonnell Center for Systems Neuroscience at Washington University.

## AUTHOR CONTRIBUTIONS

M.L.: Writing, Visualization, Validation, Software, Methodology, Investigation, Conceptualization.

## REFERENCES

1. Gonzalez-Castillo J, Bandettini PA. Task-based dynamic functional connectivity: Recent findings and open questions. NeuroImage. 2018 Oct 15;180:526–33.

2. Poline JB, Brett M. The general linear model and fMRI: Does love last forever? NeuroImage. 2012 Aug 15;62(2):871–80.

3. Bandettini PA. Twenty years of functional MRI: The science and the stories. NeuroImage. 2012 Aug 15;62(2):575–88.

4. Engel SA, Glover GH, Wandell BA. Retinotopic organization in human visual cortex and the spatial precision of functional MRI. Cereb Cortex. 1997 Mar 1;7(2):181–92.

5. Saarimäki H. Naturalistic Stimuli in Affective Neuroimaging: A Review. Front Hum Neurosci [Internet]. 2021 June 17 [cited 2025 Dec 1];15. Available from: https://www.frontiersin.org/journals/human-neuroscience/articles/10.3389/fnhum.2021.675068/full

6. Demirtaş M, Ponce-Alvarez A, Gilson M, Hagmann P, Mantini D, Betti V, et al. Distinct modes of functional connectivity induced by movie-watching. NeuroImage. 2019 Jan 1;184:335–48.

7. Vanderwal T, Kelly C, Eilbott J, Mayes LC, Castellanos FX. *Inscapes*: A movie paradigm to improve compliance in functional magnetic resonance imaging. NeuroImage. 2015 Nov 15;122:222–32.

8. Meer JN van der, Breakspear M, Chang LJ, Sonkusare S, Cocchi L. Movie viewing elicits rich and reliable brain state dynamics. Nat Commun. 2020 Oct 5;11(1):5004.

9. Kauppi JP, Jääskeläinen IP, Sams M, Tohka J. Inter-subject correlation of brain hemodynamic responses during watching a movie: localization in space and frequency. Front Neuroinform [Internet]. 2010 Mar 19 [cited 2025 Dec 1];4. Available from: https://www.frontiersin.org/journals/neuroinformatics/articles/10.3389/fninf.2010.00005/full

10. Simony E, Chang C. Analysis of stimulus-induced brain dynamics during naturalistic paradigms. NeuroImage. 2020 Aug 1;216:116461.

11. Bordier C, Puja F, Macaluso E. Sensory processing during viewing of cinematographic material: Computational modeling and functional neuroimaging. NeuroImage. 2013 Feb 15;67:213–26.

12. Russ BE, Leopold DA. Functional MRI mapping of dynamic visual features during natural viewing in the macaque. NeuroImage. 2015 Apr 1;109:84–94.

13. Nishimoto S, Vu AT, Naselaris T, Benjamini Y, Yu B, Gallant JL. Reconstructing Visual Experiences from Brain Activity Evoked by Natural Movies. Current Biology. 2011 Oct 11;21(19):1641–6.

14. Hu X, Li K, Han J, Hua X, Guo L, Liu T. Bridging the Semantic Gap via Functional Brain Imaging. IEEE Transactions on Multimedia. 2012 Apr;14(2):314–25.

15. Hu X, Deng F, Li K, Zhang T, Chen H, Jiang X, et al. Bridging low-level features and high-level semantics via fMRI brain imaging for video classification. In: Proceedings of the 18th ACM international conference on Multimedia [Internet]. New York, NY, USA: Association for Computing Machinery; 2010 [cited 2025 Dec 1]. p. 451–60. (MM ’10). Available from: https://dl.acm.org/doi/10.1145/1873951.1874016

16. Sonkusare S, Breakspear M, Guo C. Naturalistic Stimuli in Neuroscience: Critically Acclaimed. Trends in Cognitive Sciences. 2019 Aug 1;23(8):699–714.

17. Isik AI, Vessel EA. From Visual Perception to Aesthetic Appeal: Brain Responses to Aesthetically Appealing Natural Landscape Movies. Front Hum Neurosci [Internet]. 2021 July 21 [cited 2025 Dec 1];15. Available from: https://www.frontiersin.org/journals/human-neuroscience/articles/10.3389/fnhum.2021.676032/full

18. Lefèvre J, Baillet S. Optical flow approaches to the identification of brain dynamics. Hum Brain Mapp. 2009 Apr 17;30(6):1887–97.

19. Huth AG, Lee T, Nishimoto S, Bilenko NY, Vu AT, Gallant JL. Decoding the Semantic Content of Natural Movies from Human Brain Activity. Front Syst Neurosci [Internet]. 2016 Oct 7 [cited 2025 Dec 1];10. Available from: https://www.frontiersin.org/journals/systems-neuroscience/articles/10.3389/fnsys.2016.00081/full

20. Zhang Y, Han K, Worth R, Liu Z. Connecting concepts in the brain by mapping cortical representations of semantic relations. Nat Commun. 2020 Apr 20;11(1):1877.

21. Huth AG, Nishimoto S, Vu AT, Gallant JL. A Continuous Semantic Space Describes the Representation of Thousands of Object and Action Categories across the Human Brain. Neuron. 2012 Dec 20;76(6):1210–24.

22. Thye M, Hoffman P, Mirman D. The neural basis of naturalistic semantic and social cognition. Sci Rep. 2024 Mar 21;14(1):6796.

23. Rakhimberdina Z, Jodelet Q, Liu X, Murata T. Natural Image Reconstruction From fMRI Using Deep Learning: A Survey. Front Neurosci [Internet]. 2021 Dec 20 [cited 2025 Dec 1];15. Available from: https://www.frontiersin.org/journals/neuroscience/articles/10.3389/fnins.2021.795488/full

24. Yu S, Shi E, Wang R, Zhao S, Liu T, Jiang X, et al. A hybrid learning framework for fine-grained interpretation of brain spatiotemporal patterns during naturalistic functional magnetic resonance imaging. Front Hum Neurosci [Internet]. 2022 Sept 30 [cited 2025 Dec 1];16. Available from: https://www.frontiersin.org/journals/human-neuroscience/articles/10.3389/fnhum.2022.944543/full

25. Wen H, Shi J, Zhang Y, Lu KH, Cao J, Liu Z. Neural Encoding and Decoding with Deep Learning for Dynamic Natural Vision. Cereb Cortex. 2018 Dec 1;28(12):4136–60.

26. Sohn W, Di X, Liang Z, Zhang Z, Biswal BB. Explorations of using a convolutional neural network to understand brain activations during movie watching. psychoradiology. 2024 Mar 1;4:kkae021.

27. Liu X, Zhang Z, Nie J. Talking to the brain: Using Large Language Models as Proxies to Model Brain Semantic Representation [Internet]. arXiv; 2025 [cited 2025 Dec 1]. Available from: http://arxiv.org/abs/2502.18725

28. Villanueva CKT, Tu JC, Tripathy M, Lane C, Iyer R, Scotti PS. Predicting Brain Responses To Natural Movies With Multimodal LLMs [Internet]. arXiv; 2025 [cited 2025 Dec 1]. Available from: http://arxiv.org/abs/2507.19956

29. Nakagi Y, Matsuyama T, Koide-Majima N, Yamaguchi HQ, Kubo R, Nishimoto S, et al. Unveiling Multi-level and Multi-modal Semantic Representations in the Human Brain using Large Language Models [Internet]. bioRxiv; 2024 [cited 2025 Dec 1]. p. 2024.02.06.579077. Available from: https://www.biorxiv.org/content/10.1101/2024.02.06.579077v3

30. Zheng R, Sun L. LLM4Brain: Training a Large Language Model for Brain Video Understanding [Internet]. arXiv; 2024 [cited 2025 Dec 1]. Available from: http://arxiv.org/abs/2409.17987

31. Fu M, Chen G, Zhang Y, Zhang M, Wang Y. Comprehensive Neural Representations of Naturalistic Stimuli through Multimodal Deep Learning. eLife [Internet]. 2025 Aug 26 [cited 2025 Dec 1];14. Available from: https://elifesciences.org/reviewed-preprints/107607

32. Team G, Anil R, Borgeaud S, Alayrac JB, Yu J, Soricut R, et al. Gemini: A Family of Highly Capable Multimodal Models [Internet]. arXiv; 2025 [cited 2025 Dec 1]. Available from: http://arxiv.org/abs/2312.11805

33. Elam JS, Glasser MF, Harms MP, Sotiropoulos SN, Andersson JLR, Burgess GC, et al. The Human Connectome Project: A retrospective. NeuroImage. 2021 Dec 1;244:118543.

34. Van Essen DC, Smith SM, Barch DM, Behrens TEJ, Yacoub E, Ugurbil K. The WU-Minn Human Connectome Project: An overview. NeuroImage. 2013 Oct 15;80:62–79.

35. Van Essen DC, Ugurbil K, Auerbach E, Barch D, Behrens TEJ, Bucholz R, et al. The Human Connectome Project: A data acquisition perspective. NeuroImage. 2012 Oct 1;62(4):2222–31.

36. Jenkinson M, Beckmann CF, Behrens TEJ, Woolrich MW, Smith SM. FSL. NeuroImage. 2012;62(2):782–90.

37. Dale AM, Fischl B, Sereno MI. Cortical surface-based analysis. I. Segmentation and surface reconstruction. NeuroImage. 1999 Feb;9(2):179–94.

38. Robinson EC, Garcia K, Glasser MF, Chen Z, Coalson TS, Makropoulos A, et al. Multimodal surface matching with higher-order smoothness constraints. NeuroImage. 2018 Feb 15;167:453–65.

39. Andersson JLR, Skare S, Ashburner J. How to correct susceptibility distortions in spin-echo echo-planar images: application to diffusion tensor imaging. NeuroImage. 2003 Oct 1;20(2):870–88.

40. Salimi-Khorshidi G, Douaud G, Beckmann CF, Glasser MF, Griffanti L, Smith SM. Automatic denoising of functional MRI data: Combining independent component analysis and hierarchical fusion of classifiers. NeuroImage. 2014;90:449–68.

41. Glasser MF, Coalson TS, Robinson EC, Hacker CD, Harwell J, Yacoub E, et al. A multi-modal parcellation of human cerebral cortex. Nature. 2016 Aug;536(7615):171–8.

42. Cutting JE, Brunick KL, Candan A. Perceiving event dynamics and parsing Hollywood films. Journal of Experimental Psychology: Human Perception and Performance. 2012;38(6):1476– 90.

43. Rigotti M, Barak O, Warden MR, Wang XJ, Daw ND, Miller EK, et al. The importance of mixed selectivity in complex cognitive tasks. Nature. 2013 May;497(7451):585–90.

44. Fusi S, Miller EK, Rigotti M. Why neurons mix: high dimensionality for higher cognition. Current Opinion in Neurobiology. 2016 Apr 1;37:66–74.

45. Buckner RL, Krienen FM, Yeo BTT. Opportunities and limitations of intrinsic functional connectivity MRI. Nat Neurosci. 2013 July;16(7):832–7.

46. Sydnor VJ, Larsen B, Bassett DS, Alexander-Bloch A, Fair DA, Liston C, et al. Neurodevelopment of the association cortices: Patterns, mechanisms, and implications for psychopathology. Neuron. 2021 Sept 15;109(18):2820–46.

47. Vogel JW, Alexander-Bloch AF, Wagstyl K, Bertolero MA, Markello RD, Pines A, et al. Deciphering the functional specialization of whole-brain spatiomolecular gradients in the adult brain. Proc Natl Acad Sci USA. 2024 June 18;121(25):e2219137121.

48. Knodt AR, Elliott ML, Whitman ET, Winn A, Addae A, Ireland D, et al. Test–retest reliability and predictive utility of a macroscale principal functional connectivity gradient. Human Brain Mapping. 2023;44(18):6399–417.

49. Crittenden BM, Mitchell DJ, Duncan J. Recruitment of the default mode network during a demanding act of executive control. Van Essen DC, editor. eLife. 2015 Apr 13;4:e06481.

50. Buckner RL, Carroll DC. Self-projection and the brain. Trends Cogn Sci. 2007 Feb;11(2):49– 57.

51. Binder JR, Desai RH, Graves WW, Conant LL. Where is the semantic system? A critical review and meta-analysis of 120 functional neuroimaging studies. Cereb Cortex. 2009 Dec;19(12):2767–96.

52. Wang JY, Sukiennik N, Li T, Su W, Hao Q, Xu J, et al. A Survey on Human-Centric LLMs [Internet]. arXiv; 2024 [cited 2025 Dec 3]. Available from: http://arxiv.org/abs/2411.14491

53. Team G, Georgiev P, Lei VI, Burnell R, Bai L, Gulati A, et al. Gemini 1.5: Unlocking multimodal understanding across millions of tokens of context [Internet]. arXiv; 2024 [cited 2025 Dec 3]. Available from: http://arxiv.org/abs/2403.05530

54. Doerig A, Kietzmann TC, Allen E, Wu Y, Naselaris T, Kay K, et al. High-level visual representations in the human brain are aligned with large language models. Nat Mach Intell. 2025 Aug;7(8):1220–34.

55. Benara V, Singh C, Morris JX, Antonello R, Stoica I, Huth AG, et al. Crafting Interpretable Embeddings by Asking LLMs Questions [Internet]. arXiv; 2024 [cited 2025 Dec 3]. Available from: http://arxiv.org/abs/2405.16714

56. Sun F, Cui D, Jiao Q, Niu J, Zhang X, Shi Y, et al. The co-activation pattern between the DMN and other brain networks affects the cognition of older adults: evidence from naturalistic stimulation fMRI data. Cerebral Cortex. 2023 Dec 2;bhad466.

57. Liu X, Zhang N, Chang C, Duyn JH. Co-activation patterns in resting-state fMRI signals. NeuroImage. 2018 Oct 15;180:485–94.

58. Chen JE, Chang C, Greicius MD, Glover GH. Introducing co-activation pattern metrics to quantify spontaneous brain network dynamics. NeuroImage. 2015;111:476–88.

59. Parisi GI, Kemker R, Part JL, Kanan C, Wermter S. Continual Lifelong Learning with Neural Networks: A Review. Neural Networks. 2019 May;113:54–71.

60. Lake BM, Baroni M. Human-like systematic generalization through a meta-learning neural network. Nature. 2023 Nov;623(7985):115–21.

61. Pulvermüller F, Tomasello R, Henningsen-Schomers MR, Wennekers T. Biological constraints on neural network models of cognitive function. Nat Rev Neurosci. 2021 Aug;22(8):488–502.

